# Automatic inference of cell neighborhood in 2D and 3D using nuclear markers

**DOI:** 10.1101/2021.07.14.452382

**Authors:** Bruno Moretti, Santiago N. Rodriguez Alvarez, Hernán E. Grecco

## Abstract

**Significance:** Estimating neighboring cells by using only nuclear markers is crucial in many biological applications. Although several strategies have been used for this purpose, most published methods lack a rigorous characterization of their efficiencies. Remarkably, previously described methods are not automatic and depend only on cell-cell distance, neglecting the importance of pair-neighborhood interaction.

**Aim:** To develop a robust and automatic method for assessing cell local neighborhood, while analyzing the impact of the physical variables involved in this task.

**Approach:** We inferred neighbors from images with nuclei labeling by approximating the cell-cell interaction graph by the Delaunay triangulation of nuclei centroids. Each edge of this graph was filtered by thresholding in cell-cell distance and the maximum angle that each pair subtends with shared neighbors (pair-neighborhood interaction). Thresholds were calculated by maximizing a new robust statistic that measures the communicability efficiency of the cell graph. Using a variety of images of diverse tissues with additional membrane labeling to find the ground truth, we characterized the assessment performance.

**Results:** On average, our method detected 95% of true neighbors, with only 6% of false discoveries. Even though our method’s performance and tissue regularity are correlated, it works with performance metrics over 86% in very different organisms, including *Drosophila melanogaster*, *Tribolium castaneum*, *Arabidopsis thaliana* and *C. elegans*.

**Conclusions:** We automatically estimated neighboring relationships between cells in 2D and 3D using only nuclear markers. To achieve this goal, we filtered the Delaunay triangulation of nuclei centroids with a new measure of graph communicability efficiency. In addition, we found that taking pair-neighborhood interactions into account, in contrast to considering only cell-cell distances, leads to significant performance improvements. This becomes more notorious when the number of cells is low or the geometry of the cell graph is highly complex.

## 1 Introduction

Understanding the emergence of tissue organization requires quantifying the local and global signaling cues that each cell perceives from the environment. Among these cues, cell-cell interaction is one of the strongest to determine cellular fate during development and maintain homeostasis in adult life. For example, cell-cell contact has an important role during cell migration. A cell in a migrating tissue can change the direction of its movement after direct contact with another cell, in a process that is known as contact inhibition of locomotion (CIL). The number of neighboring cells can alter the outcome of CIL: in a sheet of cells with a free edge, only the cells at the free edge produce lamellipodia, which in turn induces a collective and directed migration of the whole tissue [1]. The number of neighboring cells in a tissue also plays an important role in proliferation. When normal, noncancerous cells are in crowded conditions, they cease proliferation and cell division. This characteristic is lost when cells undergo malignant transformation, resulting in uncontrolled proliferation and tumor formation [2].

At the molecular level, cell-cell interaction is also crucial in a plethora of biological processes. For example, the Notch signaling pathway is highly conserved and plays a key role in cell-cell communication. Since most of this pathway’s ligands are transmembrane proteins, signalling is restricted to neighboring cells [3]. In vertebrate embryonic development, the morphological segments—termed somites—that prefigure the bones and muscles of the adult are formed at the posterior end of the elongating body axis. A highly conserved biological clock consisting of an oscillatory gene regulatory network rhythmically differentiates cells into somites [4]. Here, the Notch signaling pathway plays a key role at synchronizing oscillations between neighboring cells [5]. Disruption of the Notch pathway during development causes severe somite malformation in zebrafish, chick and mouse embryos [6–11].

Thus, knowledge of neighbor relations between cells in tissues is crucial to understand the emergent collective behaviours that arise from individual cell-cell interactions. Assessing neighborhood in tissues using fluorescence microscopy, i.e. precisely identifying which cells share a border, requires labeling cellular membranes. While this is routinely done in a variety of experiments, it is also many times put aside for technical reasons or convenience. In contrast, nuclei labeling is ubiquitously found in literature as it is routinely used to count cells and identify seed points from which segmentation begins [12–14]. A variety of nuclear labels exist both for fixed and live-cell imaging, being DAPI and H2B-FP among the most common. Due to its brightness, nuclear labels are also used to perform autofocus routines and track cell movements over time [15–18].

In this work, we developed a fast, automatic and easy to implement method to find the first neighbors of cells in tissues using only a nuclear marker. To validate our method, we applied it to datasets containing both a nuclear and a membrane marker from a variety of organisms such as *Drosophila melanogaster*, *Tribolium castaneum*, *Arabidopsis thaliana* and *C. elegans*. These datasets were manually analyzed to generate ground truth images, which were in turn used to compare our automatically-found results against. We found that our method is robust and precise for all the test datasets we used, covering 95% of true neighboring relationships with 94% precision, on average.

## 2 Methods

### 2.1 Delaunay triangulation approximates neighboring cells

The goal of our method is to estimate which cells are in contact based only on the positions of nuclei centroids (Fig 1a-b). Considering this restriction, we define neighboring cells as those which are (1) close together and (2) not interfered by any other cell. This implies that the building blocks of the cell graph are not edges but triangles, because they are the minimal structures that can provide information about the influence of the graph over a given pair of nodes. Consequently, a triangulation of nuclei centroids could be a good approximation of the neighboring cells network (Fig. 1c). In particular, a Delaunay triangulation is outstanding for this task due to its special properties, which result from being the dual of the Voronoi diagram [19]. The latter divides a set of points *S* into touching convex regions, each of which contains only one point in *S*. If two regions are in contact in the Voronoi diagram, they are linked in the Delaunay triangulation. As a result of this, the Delaunay triangulation includes the nearest neighbor graph, which is a necessary (but not sufficient) requirement for any approximation of cell-cell physical interactions. In addition to this, given that Voronoi cells are geometrically stable with respect to small changes of the points in *S*, so the edges of the Delaunay triangulation are [20]. This property ensures that small displacements of cells do not affect the results, distinguishing this triangulation from others. Another essential feature in planar spaces is that the Delaunay triangulation maximizes the minimum angle in any triangle, avoiding sharp structures that would appear if distant cells were assumed to be connected [21].

**Fig. 1.**
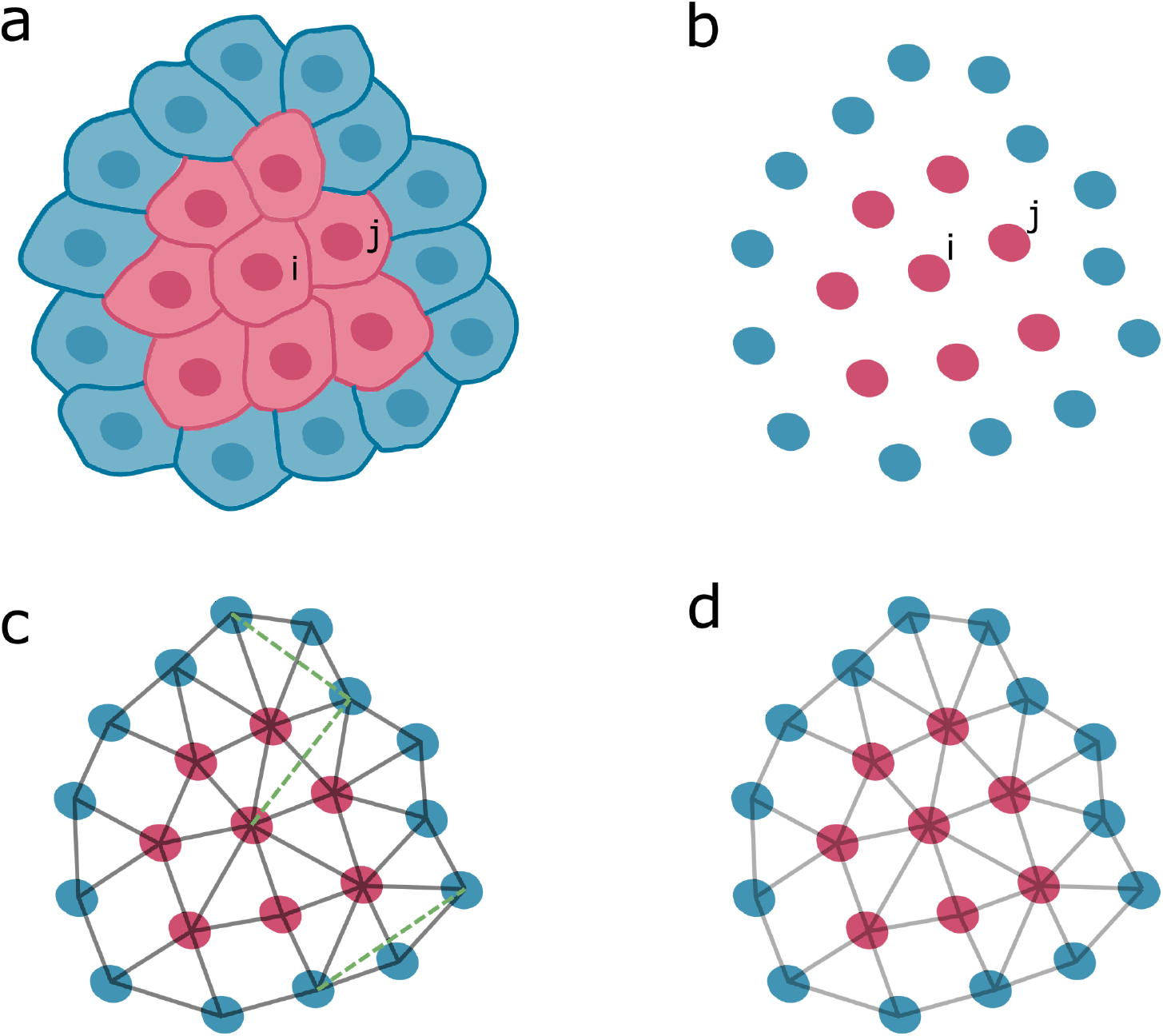
Overview of the proposed method. **(a)** System of touching cells, showing membranes and nuclei. Cell *i* and its neighbors *j* are highlighted (red). **(b)** Nuclei corresponding to the cells shown in (a). **(c)** Triangulation of nuclei centroids, including neighboring cells (gray) and typical errors (green, dashed lines). **(d)** Final result obtained by filtering the triangulation shown in (b).

### 2.2 Automatic communicablity efficiency filter

By virtue of these attributes, we designed a method to find the first neighbors of cells within tissues using only nuclear markers, based on an implementation of Delaunay triangulations present in the open-source library SciPy [22]. As mentioned before, the Delaunay triangulation is necessary but not sufficient to determine true neighboring relations and, therefore, a filtering procedure is needed (Fig. 1d). To achieve this goal, we characterized each pair of nuclei (*i*, *j*) by their distance *r*_*ij*_ and the maximum angle subtended with other nuclei Θ_*ij*_ (Fig. 2a). The latter measures the interplay between (*i*, *j*) and its neighborhood, as it informs about how other cells might be blocking the path connecting *i* and *j*.

**Fig. 2.**
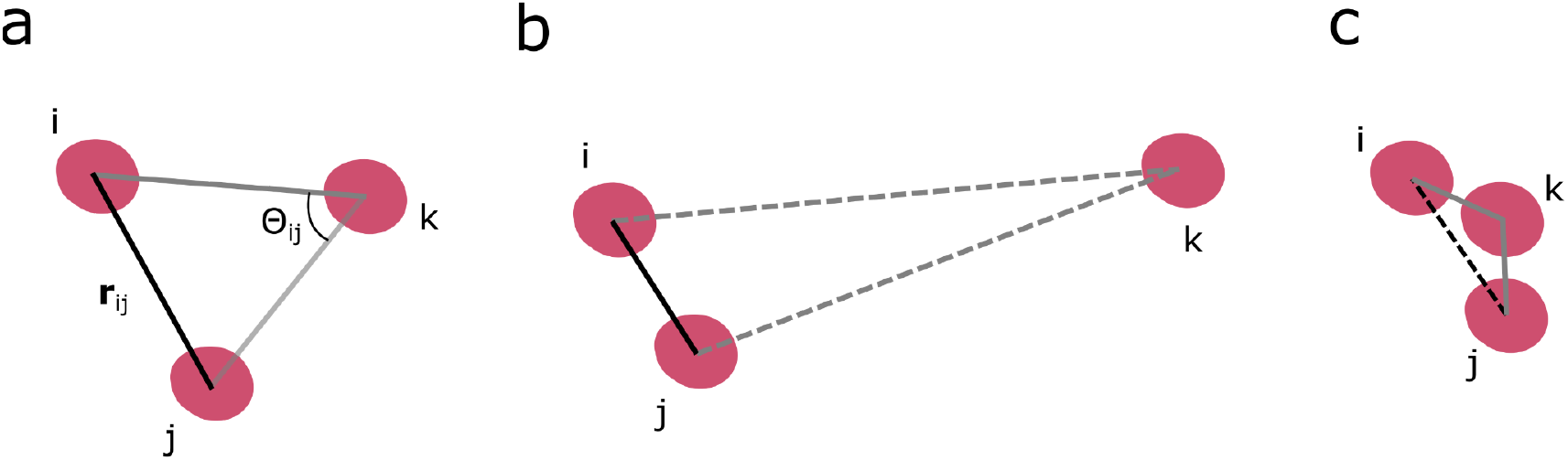
Characterization of the neighboring cells network. **(a)** Nuclei (red) are connected by solid edges representing cell membrane physical contact. Edge (*i*, *j*) is characterized by nucleus-nucleus distance *r*_*ij*_ and the maximum angle subtended with nearby nuclei Θ_*ij*_. **(b)** Small-angle limit, showing *i* and *j* close together while *k* is far away. As a result of this, (*i*, *j*) are neighbors and *k* is isolated from both *i* and *j*, which is indicated by dashed edges. **(c)** Large-angle limit, in which *k* is potentially blocking the connection between *i* and *j*.

To understand the role of Θ_*ij*_, it is instructive to analyze the possible outcomes for a set of three cells (*i*, *j, k*) in two extreme scenarios. If (*i*, *j*) are close to each other while *k* is very distant, Θ_*ij*_ is very small, indicating that (*i*, *j*) cannot be interfered by *k* (Fig. 2b.). On the other hand, when *k* is near to the midpoint of the pair, it will be very unlikely that (*i*, *j*) get in contact, no matter how close they are (Fig. 2c). Here, the relevance of Θ_*ij*_ can be appreciated: its value depends on the local spacing between cells, ranging from 0°, when (*i*, *j*) are isolated, to 180°, when there is a neighbor in between. Accordingly, (*i*, *j*) are said to be neighbors if *r*_*ij*_ ≤ *r*^*^ and Θ_*ij*_ ≤ Θ^*^, where *r* and Θ^*^ are the maximum values of distance and angle determined by the preference of the biological system towards a particular spatial organization. To estimate these parameters, and based on a previous work that introduced the concept of communicability angle between a pair of nodes in a graph [23], we developed a new statistic that measures cell-cell communicability efficiency, defined as

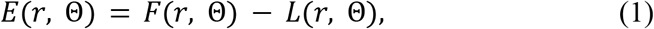

where *F* is the fraction of connected pairs and *L* is the communicability efficiency loss given that the thresholds are *r* and Θ. Consistently, the maximum communicability efficiency would be achieved by connecting the maximum number of cells at the minimum cost. The proposed critical values are those that maximize the communicability efficiency of the cell graph, i.e. thresholds are obtained by

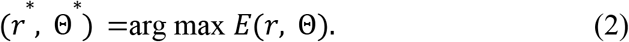

In the extreme case where there are no losses, the thresholds correspond to the minimum values of distance and angle, *r*_*min*_ and Θ_*min*_. However, this goes in detriment of the communicability efficiency, as the number of connected pairs would be at most one. Conversely, when the thresholds are *r*_*max*_ and Θ_*max*_ = 180°, then *L=1* and *F=1*, leading to null communicability efficiency. For this reason, maximizing the communicability efficiency provides a good thresholding criteria, as it balances out the benefits and costs of increasing the critical values of distance and angle.

It is worth noticing that *F* is simply calculated as the Empirical Cumulative Distribution Function (ECDF) of *r* and Θ, which corresponds to the fraction of pairs with *r*_*ij*_ ≤ *r* and Θ_*ij*_ ≤ Θ. On the other hand, *L* is not unequivocally determined by *r* and Θ. The only constraint on the loss function is that it must be an increasing function of both *r* and Θ. In this paper, we show that measuring loss as

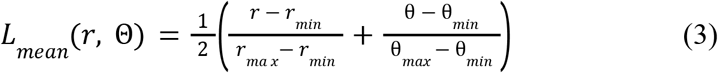

yield accurate predictions on the neighbor graph. Mathematically, *L*_*mean*_ is the mean of the normalized distance and angle. This normalization corresponds to the CDF of a random organization, in which all values of *r* and Θ are equally probable. Therefore, *L*_*mean*_ measures the communicability efficiency loss as the average probability of randomly choosing thresholds *r* and Θ .

Remarkably, our method is scale invariant and it does not depend on knowing the typical distance between neighboring cells, as this is inferred from the image itself. In the next section, we show that neighboring cells can be precisely inferred with this method in very diverse tissues.

## 3 Results

### 3.1 Delaunay triangulation-based method can be applied to diverse tissues

As a proof of concept, we applied the proposed method to images containing both nuclear and membrane fluorescent markers available in the literature. The analyzed samples included (i) *Drosophila1*: a sample of intestinal epithelium from the adult midgut of *Drosophila melanogaster* immuno-stained for the cell periphery (bPS integrin) and nuclei (DAPI) [24] (Fig. 3a), (ii) *Drosophila2*: a Drosophila embryo imaged with SiMView microscopy (Fig. 3b) [25], *Tribolium*: an average intensity projection of a uniform blastoderm of *Tribolium castaneum* embryo with H2B-RFP-labeled nuclei and GAP43-YFP-labeled membranes (Fig. 3c) [26] and *Arabidopsis*: a confocal micrograph showing the expression of different fluorescent proteins in the stem of *Arabidopsis thaliana* (Fig. 3d) [27]. The double staining present on all datasets allowed us to manually create the neighboring cells graph of each dataset, which we used to validate our estimation from only nuclei information. Edges from the estimated graph that matched the ground truth were labelled as true positive (TP). On the contrary, all pairs connected by the algorithm that were not actual neighbors were recorded as false positives (FP). Finally, false negatives (FN) were assigned to touching cells that the algorithm did not detect [28]. For details about each dataset, refer to Fig. S1 (*Drosophila1*), Fig. S2 (*Drosophila2*), Fig. S3 (*Tribolium*) and Fig. S4 (*Arabidopsis*) in the Supplemental Materials.

**Fig. 3.**
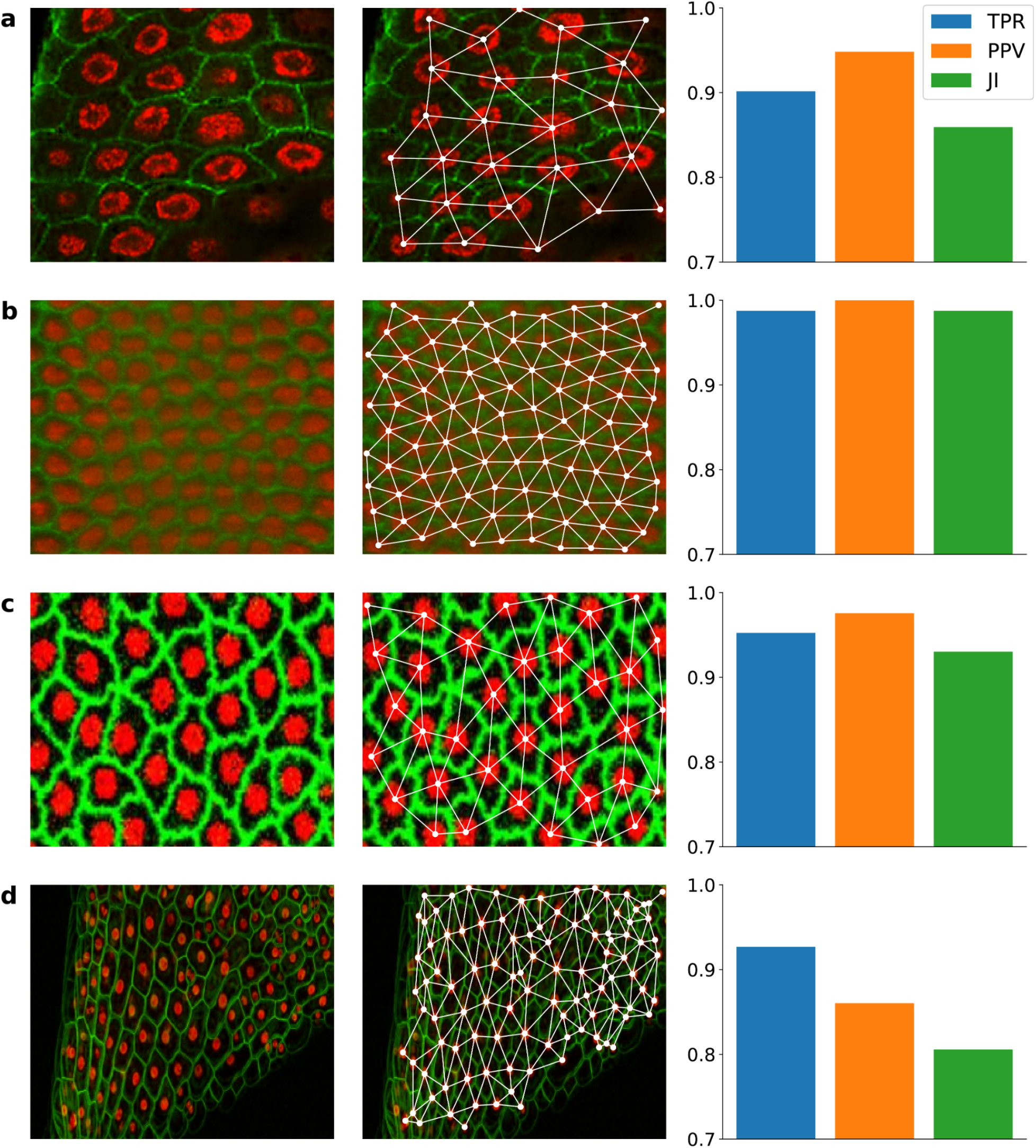
Neighboring cells estimation based on Delaunay triangulation. **(a)** Sample of intestinal epithelium from the adult midgut of *Drosophila melanogaster* immuno-stained for the cell periphery (N=27, TPR=0.902, PPV=0.948, JI=0.859). Adapted from O’Brien and Bilder 2012 [24]. **(b)** *Drosophila* embryo imaged with SiMView microscopy (N=93, TPR=0.988, PPV=1.0, JI=0.988). Adapted from Stegmaier *et al.* 2016 [25], **(c)** Average intensity projection of a uniform blastoderm of *Tribolium castaneum* embryo with H2B-RFP-labeled nuclei and GAP43-YFP-labeled membranes (N=36, TPR=0.952, PPV=0.976, JI=0.93). Adapted from Benton *et al*. 2013 [26]. **(d)** Confocal micrograph showing the expression of different fluorescent proteins in the stem of *Arabidopsis thaliana* (N=105, TPR=0.927, PPV=0.86, JI=0.806). Adapted from Federici & Haseloff 2011 [27]. **First column:** Original sample. **Second column:** filtered Delaunay triangulation of nuclei. **Third column:** performance metrics calculated by comparing the final result to the manually created ground truth. **TPR**: True Positive Rate. **PPV**: Positive Predictive Value. **JI**: Jaccard Index. Scale is not shown for images as the method is scale invariant.

Given that only a few of all possible pairs are neighbors, there are many more negatives than positives. Accordingly, performance evaluation was carried out using metrics that are focused on positives, as true negative (TN)-dependent metrics are highly biased when negatives are dominant [28]. In particular, we analyzed the True Positive Rate (TPR, also called sensitivity), Positive Predictive Value (PPV or precision) and Jaccard Index (JI). TPR measures the coverage of positives, while PPV indicates the fraction of true neighbors among all predictions. Finally, JI reports the overlap between the method and the ground truth. None of these metrics depend on the number of true negatives (TN), as they were omitted because of the imbalanced data scenario.

As it is shown in Fig. 3, TPR was over 90% in all cases, supporting the idea that the Delaunay triangulation of nuclei provides a good estimation of neighboring cells. In addition to this, PPV was 94.6% on average and a maximum of 100% was reached in a *Drosophila* embryo tissue with highly regular spatial distribution (Fig. 3b). This indicates that the more regular the tissue, the better the method’s performance. However, its range of application has proven wider as results were satisfactory in irregular tissues, such as the one from the *Arabidopsis* dataset (Fig. 3d). Even in this case, the poorest performance was 80% in JI.

### 3.2 Our method is easily extensible to 3D datasets

As Delaunay triangulations exist in all n-dimensional spaces, the proposed algorithm can be easily extended to 3D stacks. In contrast to the planar space, the basic structure of the 3D Delaunay network is not a triangle but a tetrahedron. Therefore, before applying the proposed method, each tetrahedron is decomposed in four triangles. Following steps in the method’s pipeline have no differences with those described in previous sections. To show the usefulness of this pipeline, we estimated the neighboring cells of a 3D stack of a *Caenorhabditis elegans* embryo (Fig. 4a and Video S1), whose nuclei and membranes were stained with mCherry and GFP, respectively [29]. As shown in Fig. 4b, our method was successful in all performance metrics (≥ 90%), with a remarkable TPR of 97.6%.

**Fig. 4.**
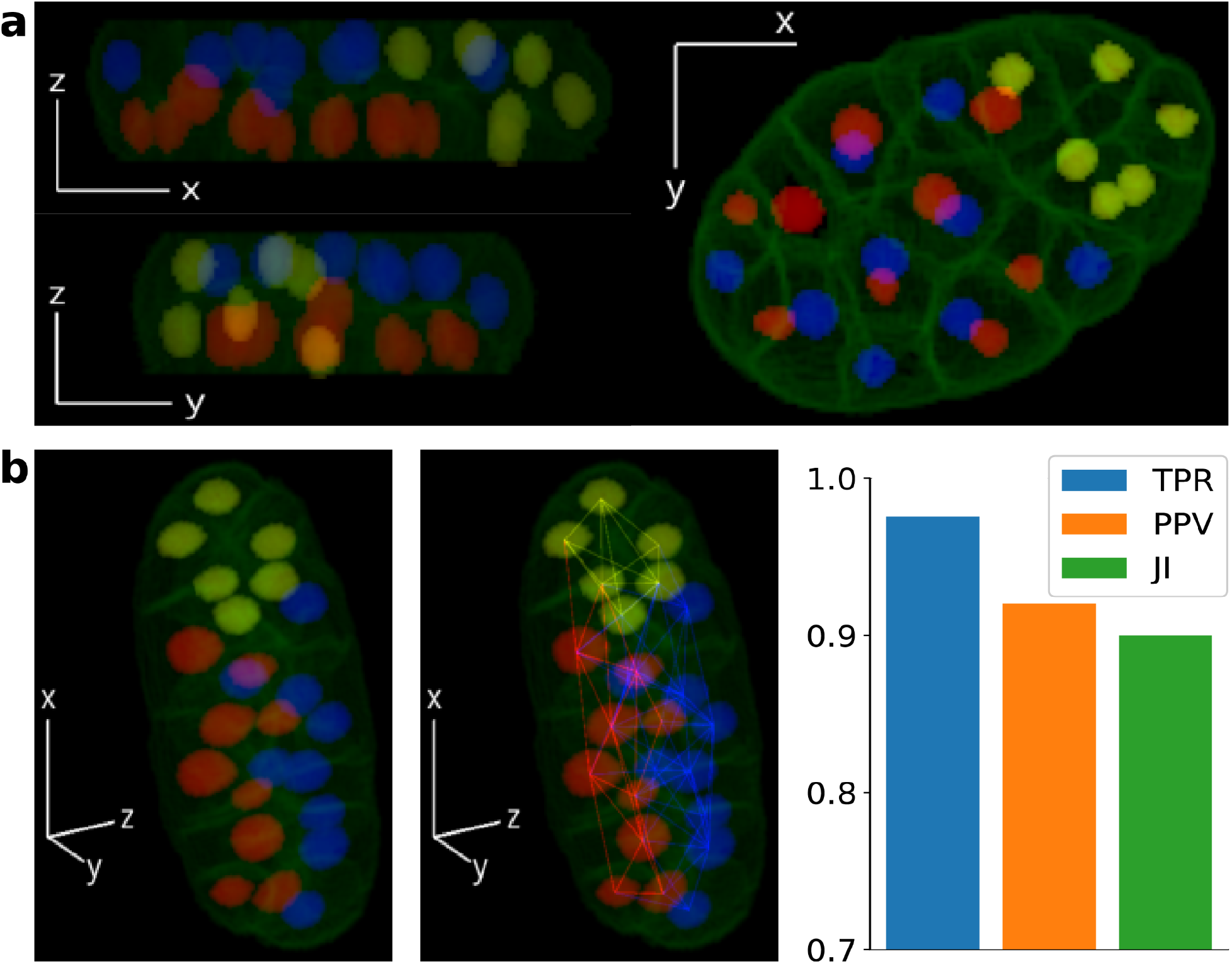
Proposed method’s results in 3D. **(a)** Planar views of a 3D stack of a *C. elegans* embryo, showing cell membranes (green) and nuclei (N=24). Colors were chosen so that the embryo was divided in *z* =8.83μm (red: *z* < 8.83μm; blue: *z* ≥8.83μm). Detected neighbors of the cell with the maximum x-position are highlighted (yellow). Adapted from Azuma and Onami 2017 [29]. **(b)** Final results: 3D view of the analyzed specimen (left), estimated cell graph (center) and performance metrics (right; TPR=0.976, PPV=0.92, JI=0.9). **TPR**: True Positive Rate. **PPV**: Positive Predictive Value. **JI**: Jaccard Index. Scale is not shown for images as the method is scale invariant.

## 4 Discussion

We developed an automatic and robust method for estimating neighboring relationships between cells using only the information provided by nuclear markers. This is a remarkable feature because membrane staining is not always possible nor convenient in many applications, ranging from cell signalling to tissue development. The robustness of our method is mostly determined by making an educated guess of neighbor relationships between cells via a Delaunay triangulation, which makes results stable against small displacements of cells. In particular, this property is useful in cell tracking, as it is expected that small fluctuations in the positions of cells between timepoints do not affect their connection. Regardless of being designed for nuclear markers, it is worth noticing that this method can also be applied to cell centroids whenever cytosolic or cytoplasmic fluorescent markers are available.

Although similar strategies have been used before [15,30,31], previous works were not focused on neighboring relationships detection and, therefore, didn’t analyze the performance of the methods directly. In addition, Delaunay edges were only filtered by distance, omitting potential interferences in cell-cell interactions that come from the presence of other cells in the neighborhood [31]. Furthermore, the distance threshold was an input of the algorithm, so that both its reusability and generalization were user-dependent. To our knowledge, we developed the first assessment of cell neighborhoods that is (i) fully automatic and (ii) based not only on cell-cell distances, but also on pair-neighborhood interactions. The latter were quantified by measuring the angle that each pair subtended with shared neighbors. The automation of our method was achieved by maximizing a new robust statistic that assigns a communicability efficiency to each edge of the Delaunay triangulation (Fig. S5).

On the other hand, incorporating relevant biological structure to the model improved the performance, as it can be seen by comparing our method’s results to those obtained by filtering the Delaunay graph only by distance (Fig. S6). Remarkably, this contribution improved PPV by +11.5% in *Drosophila1*. On average, JI and PPV were increased by +4.2% and +5.6%, respectively, while TPR did not change significantly (−0.09%). Performance differences seem to be more noticeable when the image contains a low number of cells (*Drosophila1*) or the distribution of cells is more complex (*C. elegans*). These aspects could be relevant, for instance, when tracking cell-cell interactions during early stages of embryonic development in 3D.

Importantly, we observed that tissue regularity affects the performance of our method. This may be due to the correspondence between the Delaunay triangulation and the Voronoi diagram. Since Voronoi cells are convex polygons, neighboring relations betweens highly irregular cells, such as dendritic cells or neurons, might not be well approximated by the Delaunay triangulation. A solution to this problem could be replacing nuclei centroids by a more representative set of points, determined by non-convex structures [32]. This analysis is beyond the scope of this work and could be the purpose of future research. However, the validation of our method showed that it is highly accurate and versatile, working with similar performance in diverse tissues both in 2D and 3D.

## Code, Data, and Materials Availability

Code used in this manuscript can be made available by reasonable request to the corresponding authors. Image datasets can be found in references [24–27].

---

**Fig. S1** Results obtained for the dataset *Drosophila1*.

**Fig. S2** Results obtained for the dataset *Drosophila2*.

**Fig. S3** Results obtained for the dataset *Tribolium*.

**Fig. S4** Results obtained for the dataset *Arabidopsis*.

**Fig. S5** Estimation of efficiency for all datasets

**Fig. S6** Metrics differences between the proposed method and a distance filter.

**Video S1** Animation of *C. elegans* embryo showing the estimated neighboring cell graph.

## References

1. Mayor R, Carmona-Fontaine C. Keeping in touch with contact inhibition of locomotion. Trends Cell Biol. 2010;20: 319–328.

2. Pavel M, Renna M, Park SJ, Menzies FM, Ricketts T, Füllgrabe J, et al. Contact inhibition controls cell survival and proliferation via YAP/TAZ-autophagy axis. Nat Commun. 2018;9: 2961.

3. Bray SJ. Notch signalling: a simple pathway becomes complex. Nat Rev Mol Cell Biol. 2006;7: 678–689.

4. Krol AJ, Roellig D, Dequéant M-L, Tassy O, Glynn E, Hattem G, et al. Evolutionary plasticity of segmentation clock networks. Development. 2011;138: 2783–2792.

5. Venzin OF, Oates AC. What are you synching about? Emerging complexity of Notch signaling in the segmentation clock. Dev Biol. 2020;460: 40–54.

6. Oates AC, Mueller C, Ho RK. Cooperative function of deltaC and her7 in anterior segment formation. Dev Biol. 2005;280: 133–149.

7. van Eeden FJ, Granato M, Schach U, Brand M, Furutani-Seiki M, Haffter P, et al. Mutations affecting somite formation and patterning in the zebrafish, Danio rerio. Development. 1996;123: 153–164.

8. Jiang YJ, Aerne BL, Smithers L, Haddon C, Ish-Horowicz D, Lewis J. Notch signalling and the synchronization of the somite segmentation clock. Nature. 2000;408: 475–479.

9. Dale JK, Maroto M, Dequeant M-L, Malapert P, McGrew M, Pourquie O. Periodic notch inhibition by lunatic fringe underlies the chick segmentation clock. Nature. 2003;421: 275–278.

10. Oka C, Nakano T, Wakeham A, de la Pompa JL, Mori C, Sakai T, et al. Disruption of the mouse RBP-J kappa gene results in early embryonic death. Development. 1995. pp. 3291–3301. doi:10.1242/dev.121.10.3291

11. Kusumi K, Sun ES, Kerrebrock AW, Bronson RT, Chi DC, Bulotsky MS, et al. The mouse pudgy mutation disrupts Delta homologue Dll3 and initiation of early somite boundaries. Nat Genet. 1998;19: 274–278.

12. Stanoev A, Mhamane A, Schuermann KC, Grecco HE, Stallaert W, Baumdick M, et al. Interdependence between EGFR and Phosphatases Spatially Established by Vesicular Dynamics Generates a Growth Factor Sensing and Responding Network. Cell Syst. 2018;7: 295–309.e11.

13. McQuin C, Goodman A, Chernyshev V, Kamentsky L, Cimini BA, Karhohs KW, et al. CellProfiler 3.0: Next-generation image processing for biology. PLOS Biology. 2018. p. e2005970. doi:10.1371/journal.pbio.2005970

14. Delaune EA, François P, Shih NP, Amacher SL. Single-Cell-Resolution Imaging of the Impact of Notch Signaling and Mitosis on Segmentation Clock Dynamics. Dev Cell. 2012;23: 995–1005.

15. Cervino AS, Moretti B, Stuckenholz C, Grecco HE, Davidson LA, Cecilia Cirio M. Furry is required for cell movements during gastrulation and functionally interacts with NDR1. Sci Rep. 2021;11: 1–17.

16. Wolff C, Tinevez JY, Pietzsch T, Stamataki E, Harich B, Guignard L, et al. Multi-view light-sheet imaging and tracking with the MaMuT software reveals the cell lineage of a direct developing arthropod limb. Elife. 2018;7: 1–31.

17. Attardi A, Fulton T, Florescu M, Shah G, Muresan L, Lenz MO, et al. Neuromesodermal progenitors are a conserved source of spinal cord with divergent growth dynamics. Development. 2019;146: dev175620.

18. Tomer R, Khairy K, Amat F, Keller PJ. Quantitative high-speed imaging of entire developing embryos with simultaneous multiview light-sheet microscopy. Nat Methods. 2012;9: 755–763.

19. Delaunay B. Sur la sphère vide. A la mémoire de Georges Voronoï. Bulletin de l’Académie des Sciences de l’URSS Classe des sciences mathématiques et na. 1934; 793–800.

20. Musin OR. Properties of the Delaunay triangulation. Proceedings of the thirteenth annual symposium on Computational geometry. New York, NY, USA: Association for Computing Machinery; 1997. pp. 424–426.

21. Reem D. The Geometric Stability of Voronoi Diagrams with Respect to Small Changes of the Sites. Proceedings of the twenty-seventh annual symposium on Computational geometry. New York, NY, USA: Association for Computing Machinery; 2011. pp. 254–263.

22. Virtanen P, Gommers R, Oliphant TE, Haberland M, Reddy T, Cournapeau D, et al. SciPy 1.0: fundamental algorithms for scientific computing in Python. Nat Methods. 2020;17: 261–272.

23. Estrada E, Hatano N. Communicability Angle and the Spatial Efficiency of Networks. SIAM Rev. 2016;58: 692–715.

24. O’Brien L, Bilder D. CIL:39686, Drosophila melanogaster, intestinal epithelial cell, stem cell. CIL. Dataset. 2012. doi:10.7295/W9CIL39686

25. Stegmaier J, Amat F, Lemon WC, McDole K, Wan Y, Teodoro G, et al. Real-Time Three-Dimensional Cell Segmentation in Large-Scale Microscopy Data of Developing Embryos. Dev Cell. 2016;36: 225–240.

26. Benton MA, Akam M, Pavlopoulos A. Cell and tissue dynamics during Tribolium embryogenesis revealed by versatile fluorescence labeling approaches. Development. 2013;140: 3210–3220.

27. Federici F, Haseloff J. CIL:38805, Arabidopsis thaliana. CIL. Dataset. 2011. doi:doi:10.7295/W9CIL38805

28. Luque A, Carrasco A, Martín A, de las Heras A. The impact of class imbalance in classification performance metrics based on the binary confusion matrix. Pattern Recognit. 2019;91: 216–231.

29. Azuma Y, Onami S. Biologically constrained optimization based cell membrane segmentation in C. elegans embryos. BMC Bioinformatics. 2017;18: 307.

30. Cilla R, Mechery V, Hernandez de Madrid B, Del Signore S, Dotu I, Hatini V. Segmentation and tracking of adherens junctions in 3D for the analysis of epithelial tissue morphogenesis. PLoS Comput Biol. 2015;11: e1004124.

31. Javed S, Mahmood A, Fraz MM, Koohbanani NA, Benes K, Tsang Y-W, et al. Cellular community detection for tissue phenotyping in colorectal cancer histology images. Med Image Anal. 2020;63: 101696.

32. Meyer-Hermann M. Delaunay-Object-Dynamics: cell mechanics with a 3D kinetic and dynamic weighted Delaunay-triangulation. Curr Top Dev Biol. 2008;81: 373–399.

