## Supplementary material for "Automatic inference of cell neighborhood in 2D and 3D using nuclear markers"

**Bruno Moretti,[a,b,†,*] Santiago N. Rodriguez Alvarez[a], Hernán E. Grecco[a,b,*]**
[a] Department of Physics, FCEN, University of Buenos Aires, Buenos Aires, Argentina
[b] Instituto de Física de Buenos Aires, CONICET, Buenos Aires, Argentina
† Current affiliation: Department of Molecular and Cell Biology, University of California at Berkeley

### Appendix A: Supplemental Material

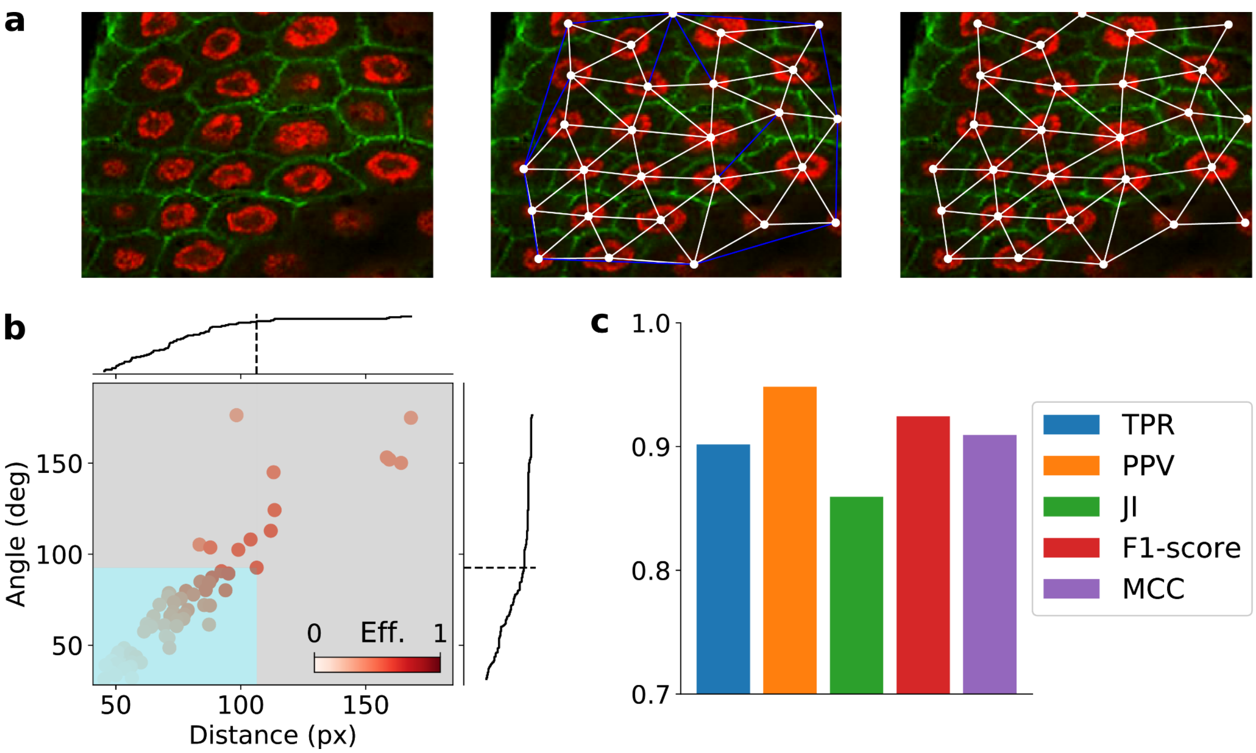


**Fig. S1** Results obtained for the dataset *Drosophila1*. **(a)** Delaunay triangulation of nuclei centroids and edge filtering by communicability efficiency. **Left**: Original sample. **Center**: Delaunay triangulation, including errors (blue). **Right:** filtered neighboring cell graph. **(b)** Distribution of Delaunay edges in the distance-angle plane. Each point represents a unique edge in the Delaunay triangulation and is colored by communicability efficiency. Coordinates of the point with maximum communicability efficiency correspond to the threshold values of distance and angle, which divide the plane in positives (cyan) and negatives (gray). **(c)** Performance metrics calculated by comparing the estimated graph to the manual ground truth. **TPR**: True Positive Rate. **PPV**: Positive Predictive Value. **JI**: Jaccard Index. **MCC:** Matthews correlation coefficient. Scale is not shown for images as the method is scale invariant.

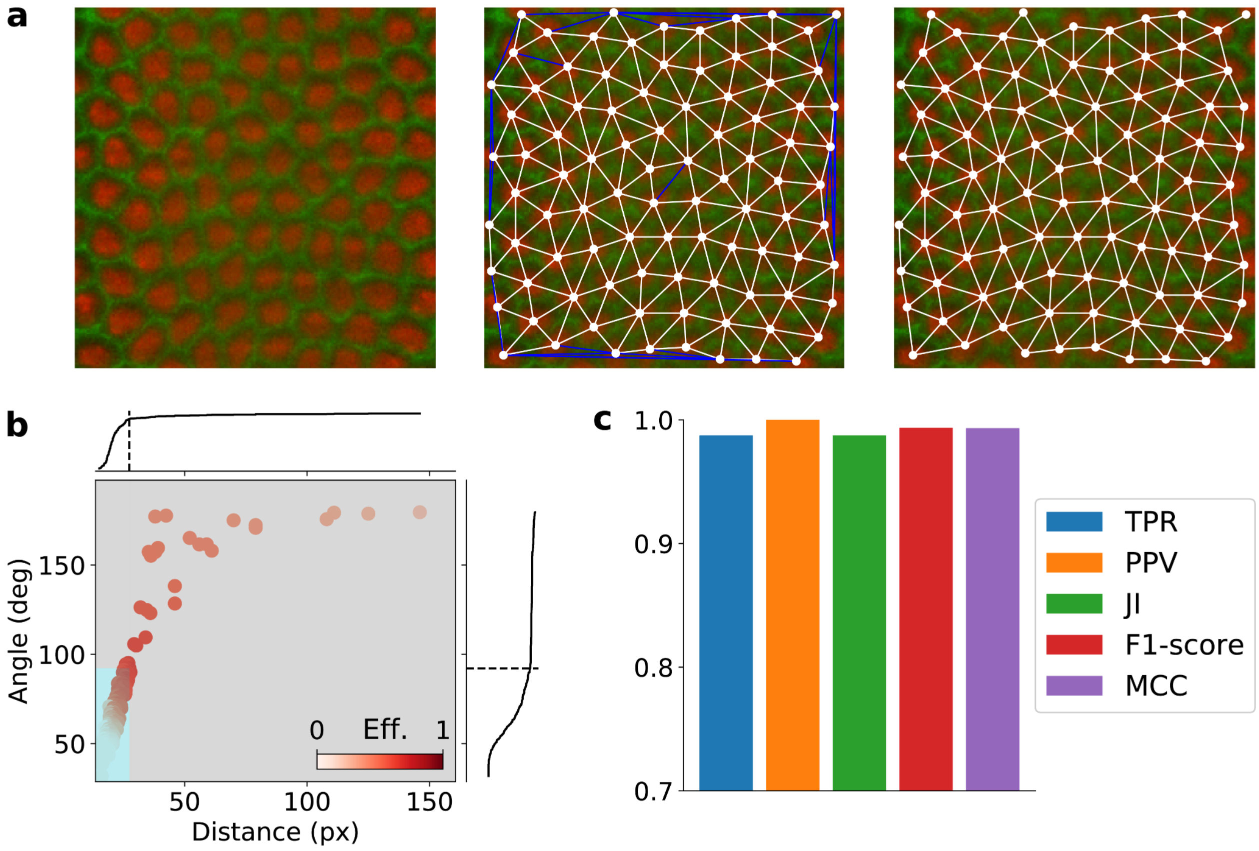


**Fig. S2** Results obtained for the dataset *Drosophila2.* **(a)** Delaunay triangulation of nuclei centroids and edge filtering by communicability efficiency. **Left**: Original sample. **Center**: Delaunay triangulation, including errors (blue). **Right:** filtered neighboring cell graph. **(b)** Distribution of Delaunay edges in the distance-angle plane. Each point represents a unique edge in the Delaunay triangulation and is colored by communicability efficiency. Coordinates of the point with maximum communicability efficiency correspond to the threshold values of distance and angle, which divide the plane in positives (cyan) and negatives (gray). **(c)** Performance metrics calculated by comparing the estimated graph to the manual ground truth. **TPR**: True Positive Rate. **PPV**: Positive Predictive Value. **JI**: Jaccard Index. **MCC:** Matthews correlation coefficient. Scale is not shown for images as the method is scale invariant.

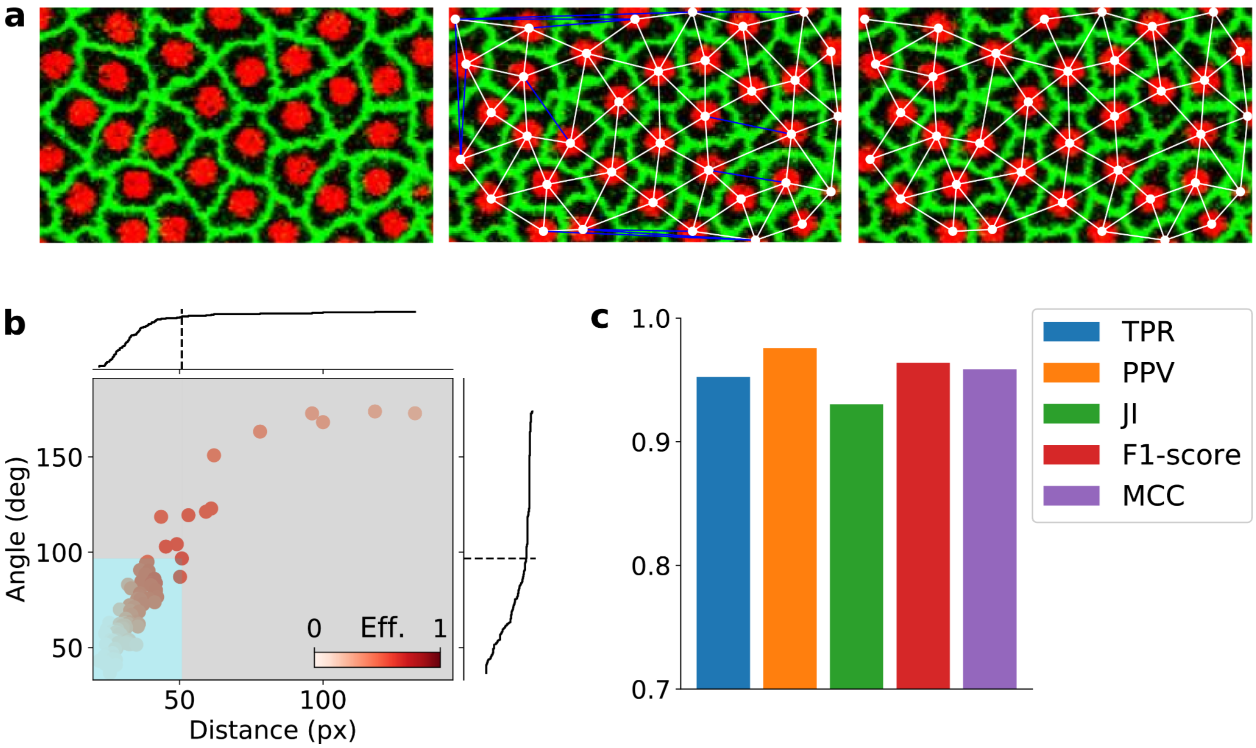


**Fig. S3** Results obtained for the dataset *Tribolium.* **(a)** Delaunay triangulation of nuclei centroids and edge filtering by communicability efficiency. **Left**: Original sample. **Center**: Delaunay triangulation, including errors (blue). **Right:** filtered neighboring cell graph. **(b)** Distribution of Delaunay edges in the distance-angle plane. Each point represents a unique edge in the Delaunay triangulation and is colored by communicability efficiency. Coordinates of the point with maximum communicability efficiency correspond to the threshold values of distance and angle, which divide the plane in positives (cyan) and negatives (gray). **(c)** Performance metrics calculated by comparing the estimated graph to the manual ground truth. **TPR**: True Positive Rate. **PPV**: Positive Predictive Value. **JI**: Jaccard Index. **MCC:** Matthews correlation coefficient. Scale is not shown for images as the method is scale invariant.

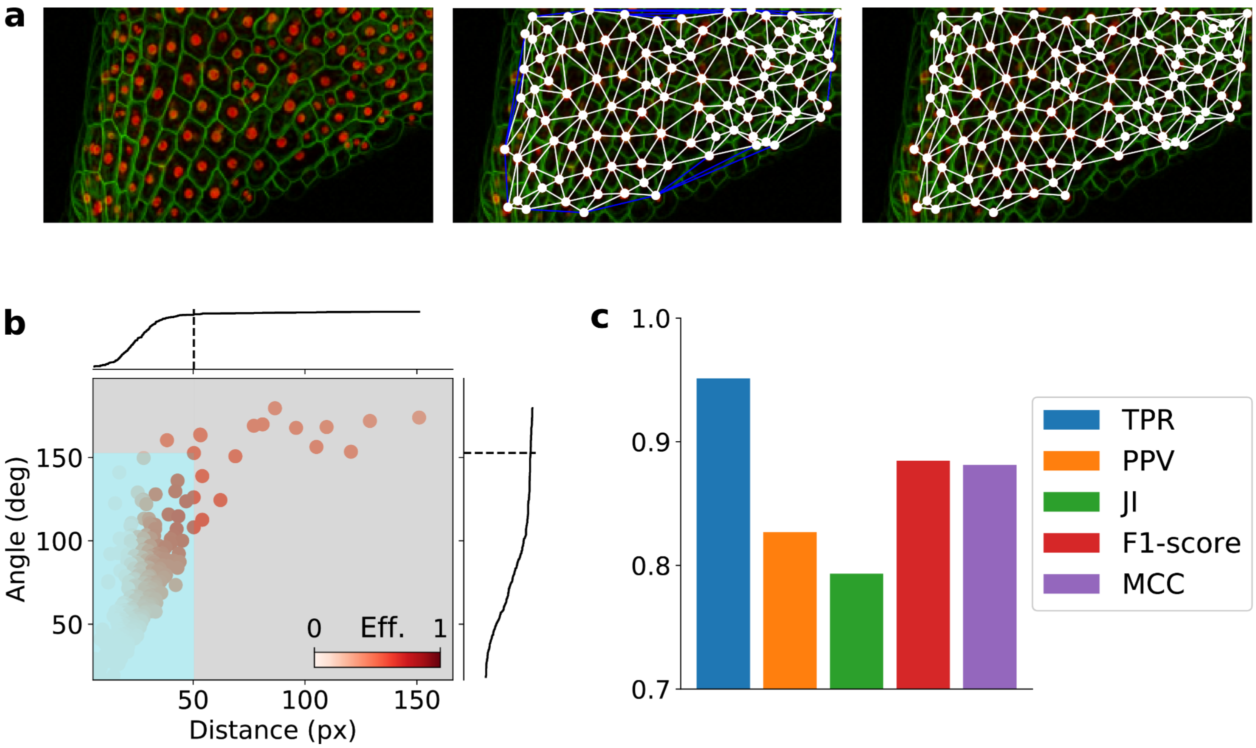


**Fig. S4** Results obtained for the dataset *Arabidopsis*. **(a)** Delaunay triangulation of nuclei centroids and edge filtering by communicability efficiency. **Left**: Original sample. **Center**: Delaunay triangulation, including errors (blue). **Right:** filtered neighboring cell graph. **(b)** Distribution of Delaunay edges in the distance-angle plane. Each point represents a unique edge in the Delaunay triangulation and is colored by communicability efficiency. Coordinates of the point with maximum communicability efficiency correspond to the threshold values of distance and angle, which divide the plane in positives (cyan) and negatives (gray). **(c)** Performance metrics calculated by comparing the estimated graph to the manual ground truth. **TPR**: True Positive Rate. **PPV**: Positive Predictive Value. **JI**: Jaccard Index. **MCC:** Matthews correlation coefficient. Scale is not shown for images as the method is scale invariant.

**Video S1** Animation of *C. elegans* embryo showing the estimated neighboring cell graph. Colors blue and red divide the embryo in two halves in Z, while yellow correspond to a particular cell at the border of the organism and its neighbors.

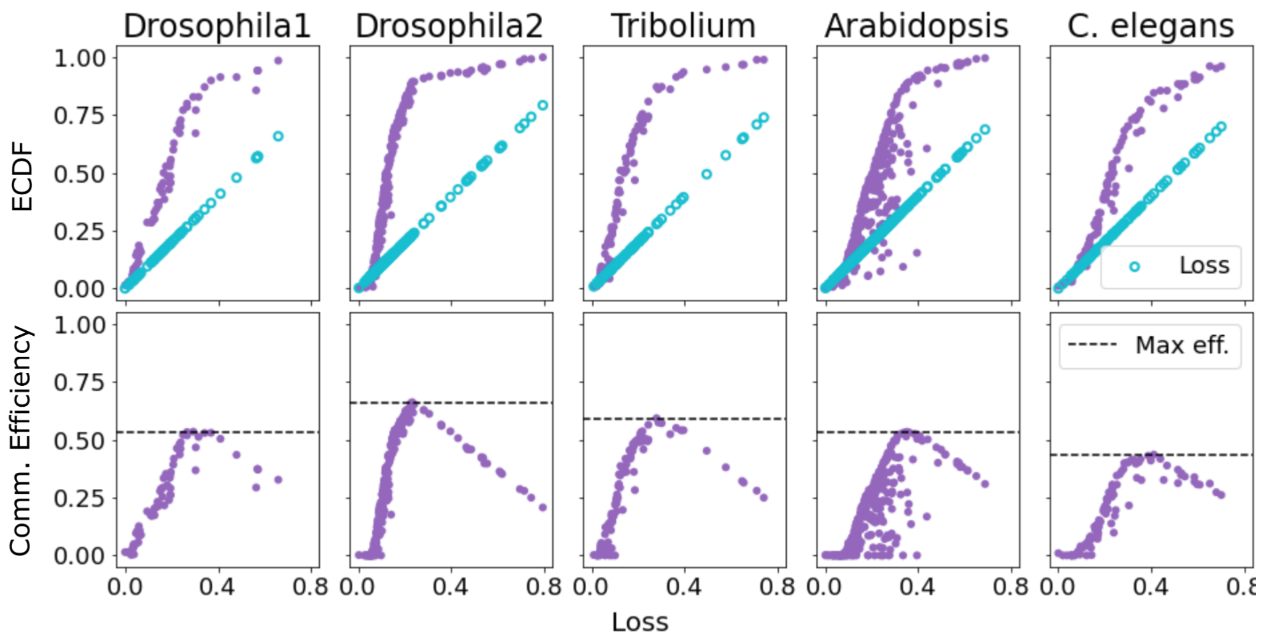


**Fig. S5** Estimation of communicability efficiency for all datasets. Points correspond to edges in the Delaunay triangulation. **First row:** Empirical Cumulative Distribution Function (ECDF) versus mean loss. To provide a reference, the mean loss function (identity) is shown. **Second row:** Communicability efficiency obtained by subtracting the mean loss to the ECDF. Maximum communicability efficiency is highlighted.

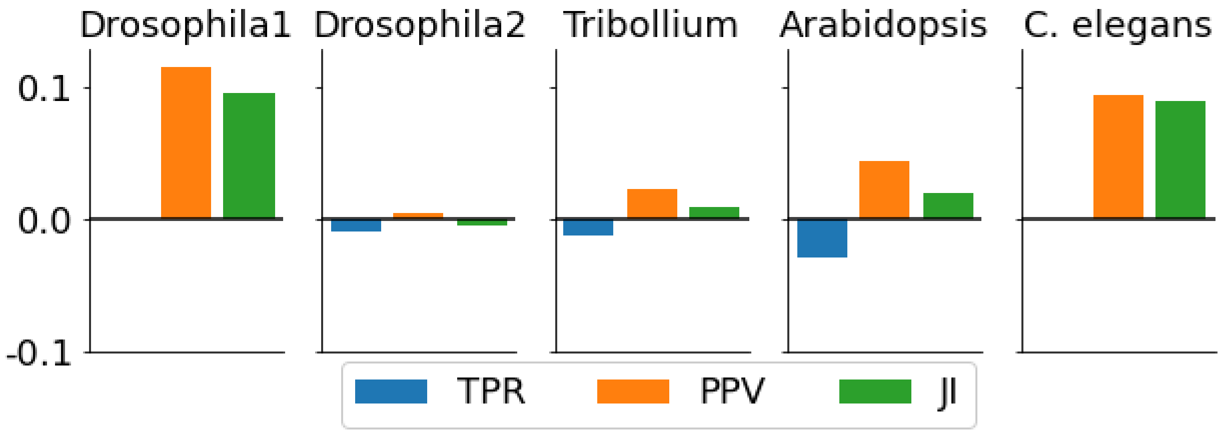


**Fig. S6** Metrics differences between our method and other methods that use only distance filters. The figure shows that improvements would become more significant with a low number of cells (*Drosophila1*) or a complex geometry (*C. elegans*). Maximum difference observed corresponds to *Drosophila1,* with a PPV advantage of +11.5%. **TPR**: True Positive Rate. **PPV**: Positive Predictive Value. **JI**: Jaccard Index.
